# Structural transitions in the GTP cap visualized by cryo-EM of catalytically inactive microtubules

**DOI:** 10.1101/2021.08.13.456308

**Authors:** Benjamin J LaFrance, Johanna Roostalu, Gil Henkin, Basil J Greber, Rui Zhang, Davide Normanno, Chloe McCollum, Thomas Surrey, Eva Nogales

**Affiliations:** Department of Molecular and Cell Biology, University of California, Berkeley, California 94720, USA; The Francis Crick Institute, London, United Kingdom; Centre for Genomic Regulation, Barcelona Institute of Science and Technology, Barcelona, Spain; California Institute for Quantitative Biosciences (QB3), University of California, Berkeley, California 94720, USA; Molecular Biophysics and Integrative Bio-Imaging Division, Lawrence Berkeley National Laboratory, Berkeley, California 94720, USA; ICREA, Barcelona, Spain; Howard Hughes Medical Institute, University of California, Berkeley, California 94720, USA

**Keywords:** microtubules, cryo-EM, TIRF, GTP, dynamic instability

## Abstract

Microtubules (MTs) are polymers of α/β-tubulin heterodimers that stochastically switch between growth and shrinkage phases. This dynamic instability is critically important for MT function. It is believed that GTP hydrolysis within the MT lattice is accompanied by destabilizing conformational changes, and that MT stability depends on a transiently existing GTP cap at the growing MT end. Here we use cryo-EM and TIRF microscopy of GTP hydrolysis-deficient MTs assembled from mutant recombinant human tubulin to investigate the structure of a GTP-bound MT lattice. We find that the GTP-MT lattice of two mutants in which the catalytically active glutamate in α-tubulin was substituted by inactive amino acids (E254A and E254N) is remarkably plastic. Undecorated E254A and E254N MTs with 13 protofilaments both have an expanded lattice, but display opposite protofilament twists, making these lattices distinct from the compacted lattice of wildtype GDP-MTs. End binding proteins of the EB family have the ability to compact both mutant GTP-lattices and to stabilize a negative twist, suggesting that they promote this transition also in the GTP cap of wildtype MTs, thereby contributing to the maturation of the MT structure. We also find that the MT seam appears to be stabilized in mutant GTP-MTs and destabilized in GDP-MTs, supporting the proposal that the seam plays an important role in MT stability. Together, these first high-resolution structures of truly GTP-bound MTs add mechanistic insight to our understanding of MT dynamic instability.

**Significance Statement:** Microtubules (MTs) are non-equilibrium polymers that switch between states of growth and shrinkage. This property is critical for their function and is a consequence of GTP hydrolysis in the MT. The structure of the stable GTP part of the MT (the GTP cap) has previously been inferred from MTs polymerized with non-hydrolyzable GTP analogs. Here, we report the first high-resolution structures of MTs truly containing GTP, polymerized from mutated, hydrolysis-deficient tubulins. We find that GTP-MTs have an “expanded lattice” and a “closed seam”, structural characteristics possibly responsible for stabilizing the GTP cap. These results provide new insight into the structural transitions at growing MT ends, furthering our understanding of the bistable nature of MTs.

## Introduction

Microtubules (MTs) form by self-assembly of GTP-bound α/β-tubulin heterodimers into a cylinder consisting of laterally associated linear protofilaments (pfs) (Fig. 1A). During incorporation into the MT lattice, GTP hydrolysis is triggered by the catalytic glutamate in α-tubulin (E254), which becomes juxtaposed to the GTP in β-tubulin at the longitudinal interface between dimers. GTP hydrolysis has a destabilizing effect on the MT structure and polymerization can only continue in the presence of a GTP-tubulin cap at the growing end of the MT. Stochastic loss of this cap is thought to lead to catastrophe, the switching into a depolymerization phase (1–3). The change between the stable GTP-tubulin and the meta-stable GDP-tubulin lattices is therefore at the heart of the phenomenon of MT dynamic instability, which is critical to MT function in the cell (4).

**Fig. 1.**
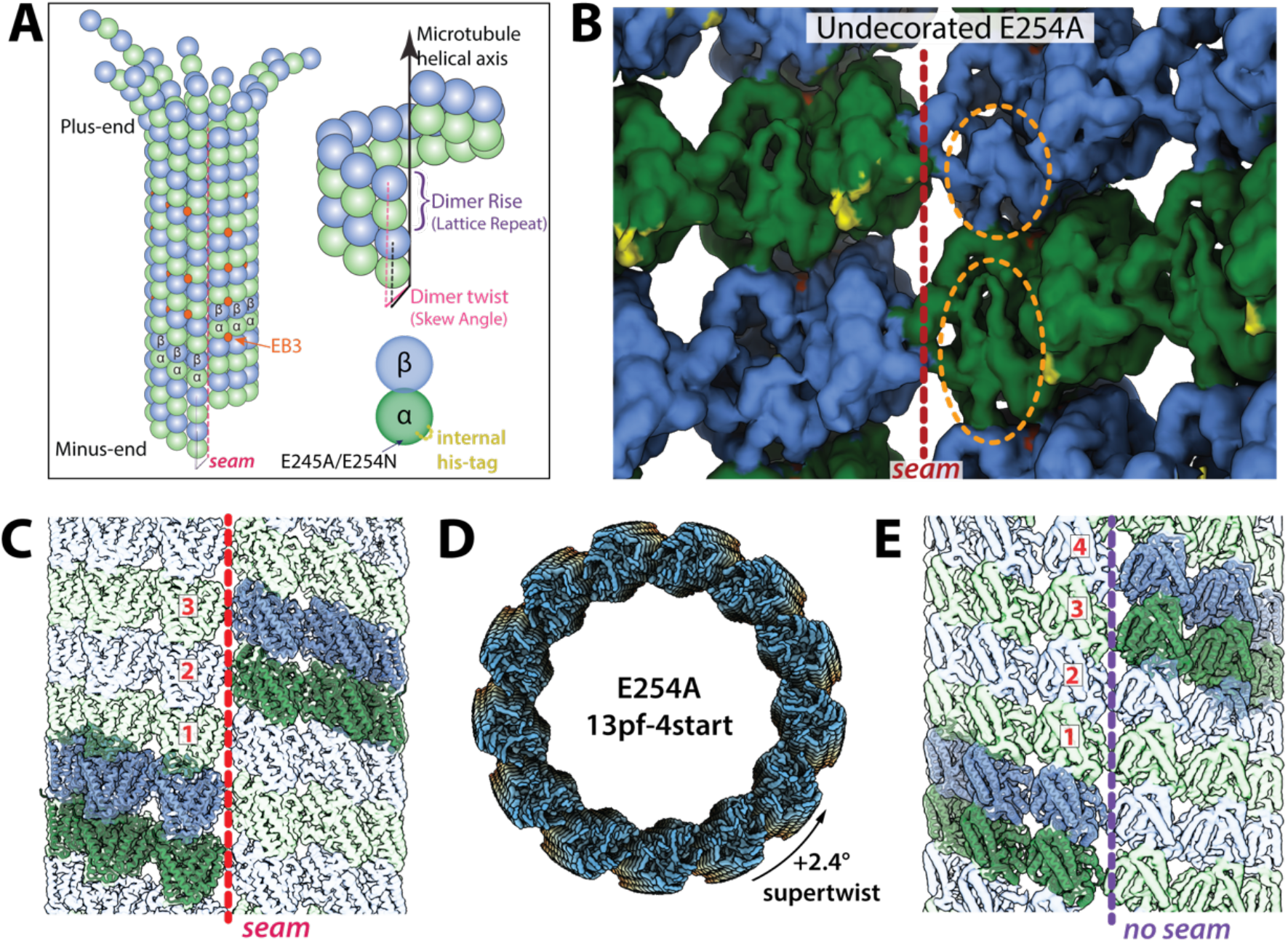
Structural characterization of E254A MTs. **(A)** Cartoon diagram of a 13pf 3-start MT undergoing depolymerization, with α-tubulin in green, β-tubulin in blue, and EB3 in orange (colors maintained throughout). Also shown are the dimer rise and dimer twist (or skew angle) of the MT lattice, where rise denotes the distance (in Å) from one tubulin dimer to the one directly above it, and twist is the angle around the MT helical axis that must occur going from one dimer to the one above. **(B)** View of tubulin dimers from the lumen, where orange ovals highlight distinctive features of the α and β tubulin subunits, and the red dashed line denotes the seam for E254A MTs. The small region highlighted in yellow corresponds to additional density in the recombinant sample that can be assigned to the internal His6-tag. **(C)** A representative 13pf 3-start MT (in this case, E254A). The same generic architecture was observed for wildtype and E254N MTs (though with subtle differences in lattice parameters). **(D)** Sub-classification of 13pf E254A dataset revealed a subset of 13pf 4-start MTs, here shown colored blue-yellow-orange along the MT-axis. **(E)**, Lateral view of 13pf 4-start MTs showing the absence of a seam in the MT lattice. In (C) and (E), one helical layer of tubulin dimers is highlighted, and the corresponding ribbon diagrams docked into those dimers, to emphasize the difference between the two lattices.

Given the critical nature of the GTP-to-GDP transition for MT dynamics, and the nucleotide-specific recruitment of specialized factors to the MT lattice, it is critical to understand the nuanced structural differences between the nucleotide states within the MT. Previous cryo-electron microscopy (cryo-EM) studies have made use of non- or slowly-hydrolyzable GTP analogs to generate surrogates of different MT lattices (5–9). However, two GTP analogs, GMPCPP and GTPγS, give rise to different MT lattices. Whereas GMPCPP-bound MTs have an “expanded”longitudinal repeat (the dimer rise) and display a slightly positive protofilament (pf) twist, GTPγS-bound MTs are compacted and display a negative pf twist, both being distinct from GDP-bound MTs that are compacted and have hardly any pf twist (9). A twist=0° would indicate that tubulin dimers are stacked in perfect vertical alignment to one another, i.e. that the pfs run parallel to the MT axis (Fig. 1A). Because GMPCPP strongly promotes MT nucleation and growth, in contrast to GTPγS, GMPCPP has often been considered the more GTP-like analog in the context of MTs. This assumption led to the model that the MT lattice is compacted upon GTP hydrolysis due to a conformational change in α-tubulin near the longitudinal interface and the nucleotide-binding pocket in β-tubulin (5, 8–10). In this proposal, the compacted GDP state is strained and contributes to the instability of the lattice. Recent studies using X-ray fiber diffraction on an increased number of GTP-analogs have suggested an alternative conformational landscape through the hydrolysis process that starts with a compacted GTP lattice that transiently expands to allow phosphate release (11). However, the structure of a GTP-bound MT lattice has not been directly visualized at high resolution. It therefore remains unclear how accurately the models derived from GTP-analogs capture the structural transitions in the real GTP cap.

Another intriguing structural feature of most MTs is a seam that breaks helical symmetry. While most lateral interactions between pfs are homotypic (*α*-*α* and β-β interactions), across the seam this interaction is heterotypic (α-β and β-α) (Fig. 1A) and has often been thought of as a possible point of weakness in the MT lattice (8, 9, 12–14). Indeed, comparison of stabilized MTs bound to GTP-analogs with GDP-MTs have shown that while the former have a circular cross section of equally distant pfs, the less stable GDP counterpart shows the two pfs at the seam breaking that cylindrical shape and being further apart (9). Whether this holds up for true GTP-MT segments is unknown. However, it raises the possibility that the seam may indeed be more stable in the GTP cap than in the GDP lattice and thereby contribute to the stabilizing effect of the cap.

In a complementary approach, fluorescence microscopy of end binding proteins of the EB family binding to dynamic MTs has been used to gain indirect insight into the nucleotide and conformational state at growing MT ends (15–22). Both in cells and in reconstituted systems, EBs transiently bind with high affinity to a region at growing MT ends that comprises hundreds of tubulins and turns over within several seconds as MTs grow, generating the characteristic ‘end tracking’ phenomenon of EBs (15, 16, 20, 23). EBs bind at the corner between four tubulins, close to where the GTP is hydrolyzed (except along the seam), suggesting that their binding affinity may be sensitive to conformational changes in the MT lattice as GTP is hydrolyzed (8, 20).

We recently showed that EBs bind strongly to hydrolysis-deficient MTs polymerized from tubulin where the catalytic glutamate in α-tubulin has been substituted by an inactive alanine (E254A) (21). Moreover, in MTs in which GTP hydrolysis is not blocked, but expected to be slowed down by a glutamate to aspartate (E254D) mutation, the EB binding region is extended, indicating that EBs indeed recognize the GTP cap (21). This also agrees with the proposal that the EB binding region at growing MT ends protects the MT from catastrophe (18, 20). However, EBs bind only very weakly to the expanded lattice of GMPCPP-MTs, the presumed mimic of GTP-MTs, whereas they bind well to the compacted lattice of GTPγS MTs proposed to resemble a post-hydrolysis state (8, 9, 20, 22). These discrepancies raised the question of whether the GTP cap really displays an expanded lattice and, if yes, what are the key structural features that ensure high affinity binding of EBs to the GTP cap but not to GMPCPP-MTs or GDP-MTs.

Here, in order to better understand the GTP conformation of MTs, we used cryo-EM to visualize MTs assembled from recombinant wildtype (wt) and mutant human tubulin in which the catalytic glutamate in α-tubulin was mutated to alanine or asparagine (E254A or E254N, respectively), making them deficient in GTP hydrolysis. We find that the catalytically inactive MTs both are stable and have an expanded dimer rise, but differ in their pf twist. E254A MTs display a negative twist and high EB-binding affinity. E254N MTs have a positive twist and show low EB binding affinity, making them similar to GMPCPP-MTs. However, in contrast to GMPCPP-MTs, growing E254N MTs can switch spontaneously into a high EB-binding conformation, a transition that is promoted by EBs. These results demonstrate that GTP-MTs can adopt at least two structurally distinct expanded states. EBs promote compaction and a negative pf twist in both mutant MTs, despite the absence of GTP hydrolysis, and the MT lattice they promote probably reflects conformational features adopted by the true GTP cap in EB decorated wildtype MTs. Lastly, catalytically inactive mutant MTs trapped in a GTP-bound state display a regularized seam, lending further support to the hypothesis that a more “closed” seam is a critical stabilizing feature of the GTP cap. Our structures provide new insights into the intrinsic regulation of MT instability and further our understanding of the pre-hydrolysis state of MTs and the preference of EBs for certain MT states.

## Results

To better understand how nucleotide state affects MT structure, we structurally characterized MTs assembled from catalytically inactive tubulin that are constitutively locked in the GTP state. Recombinant human tubulin was purified and biochemically characterized as described previously (21) (Fig. S1). We optimized our cryo-EM image analysis pipeline to deal with some unexpected challenges (Fig. S2; see below). Here we focus on the results obtained for 13-pf MTs given their abundance and physiological relevance in mammalian cells (24–26), but the characteristics we describe herein also held true for 14pf MTs (data not shown).

### Wildtype recombinant tubulin does not alter the MT architecture

We first determined the cryo-EM structure of MTs polymerized from recombinant wildtype human tubulin (Fig. S2A). Lattice parameters of these MTs, both undecorated and decorated with kinesin-1 motor domains, were nearly indistinguishable from previously reported GDP-MT structures assembled from mammalian brain tubulin (5, 8, 9) (Data Table 1). The dimer rise was 81.69Å and 81.50Å and the twist was +0.09° and +0.05, respectively, for recombinant wildtype human GDP-MTs and GDP-MTs from mammalian brain tubulin. However, there was additional density corresponding to the internal His_6_-tag located within the partly disordered acetylation loop of α-tubulin that includes lysine 40 in the recombinant structure (Fig. S2D; see also Fig. 1B). This purification tag has previously been shown to have no effect on polymerization dynamics (21, 27), and our current study confirms that it also does not alter lattice parameters compared to the endogenous wildtype mammalian MTs studied previously (Data Table 1) (27).

### E254A MTs have an expanded lattice with a negative twist

For the catalytically inactive mutant E254A (in α-tubulin), cryo-EM processing protocols developed for and applied to MTs (28, 29) failed to yield a high-resolution structure. Some of these E254A MTs had an abnormally large twist parameter, a feature that was also visible in raw images and 2D class averages (Fig. S2B) that challenged our image analysis pipeline. Further rounds of 3D classification after the initial classification by pf number (Fig. S2C) showed that there are at least two distinct lattice conformations for 13pf E254A MTs: one corresponding to the commonly observed 13pf 3-start lattice (Fig. 1B, C), and a second one with a 13pf 4-start lattice (Fig. 1D, E). The dimer rise of the 13pf 3-start structure was similar to the previously described GMPCPP-bound mammalian brain MTs, with a value of 83.18 Å and corresponding to an expanded lattice. However, the pf twist was negative (−0.22°) instead of positive (+0.23°) as observed for GMPCPP-MTs (Data Table 1). Historically, 4-start lattices have been almost exclusively observed for 15pf and 16pf MTs, and more rarely for 14pf (30). Similar to the 13pf 3-start MT, the 13pf 4-start lattice of the E254A MTs was also in the expanded conformation, with a dimer rise of 83.03 Å (Data Table 1). Furthermore, the 4-start lattice maintains the same inter-pf contacts as the 3-start lattice, with the exception that all 4-start contacts were homotypic (truly helical and without a seam) while the 13pf 3-start MTs displayed the canonical heterotypic seam (Fig. 1C, E). In conclusion, independent of the start number of the lattice, E254A MTs are always expanded, in contrast to the compacted lattice of GDP-MTs.

### E254N MTs switch between different lattice states as indicated by EB3 binding

To test if the presence of GTP rather than the E254A mutation itself has an effect on the structure of the MT lattice, we generated a more conservative E254N mutation at the catalytic site. E254N MTs grown in the presence of GTP also did not hydrolyze GTP, as demonstrated by HPLC analysis of their nucleotide content (Fig. S1). Next, we imaged mGFP-EB3 binding to unlabeled E254N MTs that grew from immobilized MT seeds by TIRF microscopy (Fig. 2A). As expected for GTP-hydrolysis deficient MTs, they grew persistently without displaying catastrophes (kymographs in Fig. 2B, C, D). However, the MTs display segments with different properties, as reflected by the long stretches of either weak or strong mGFP-EB3 signals, indicating that the lattice can be in at least two different conformations that are both stable (Fig. 2A). Occasionally the MT conformation could switch suddenly at the growing MT end from a weak to high EB3 binding state that then persisted as the MT grew (Fig. 2B). The opposite switch from growing a strongly EB3 binding lattice to a weakly EB3 binding lattice was not observed, suggesting that the high affinity conformation is preferred, but that this conformational state is hard to access when the microtubule has started growing with a low affinity lattice. The position of the boundary between lattices in the two conformations was either immobile (Fig. 2C) or could slowly move, either to the MT plus end (Fig. 2B) or the minus end (Fig. 2D), indicative of a slow and cooperative lattice transformation at the boundary either from high-to-low or from low-to-high EB3 binding affinity, respectively.

**Fig. 2.**
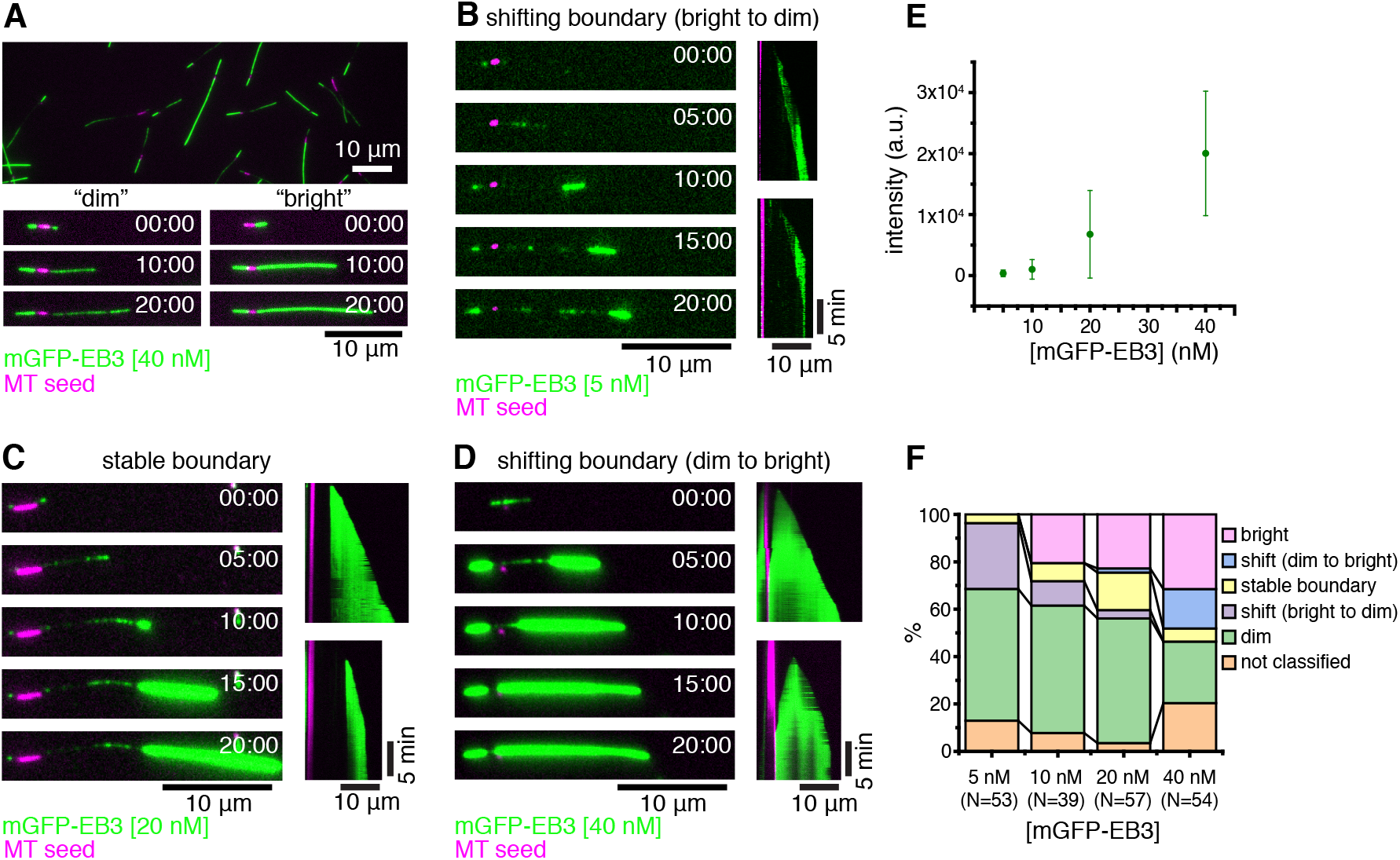
Dynamic E254N MTs observed by TIRF microscopy. **(A)** TIRF microscopy image of unlabeled non-fluorescent E254N MTs after 20 minutes of growth from CF640R-labeled GMPCPP-stabilized MT “seeds” (magenta) in the presence of 40 nM mGFP-EB3 (green) and GTP showing a variety of “dimly” and “brightly” mGFP-EB3-decorated MTs. Bottom, time series of growth of a “dim” and “bright” MT. **(B-D)** Time series and kymographs exemplifying MT growth behavior: **(B)** MT showing a switch from “dim” to “bright” lattice at the growing plus end, followed by progressive conversion of the lattice at the boundary to the “dim” state, causing the boundary to move toward the growing plus end. 5 nM mGFP-EB3. **(C)** Once this MT switches into a “bright” binding state at the growing plus end, the boundary between the “dim” and the “bright” lattice does not change. 20 nM mGFP-EB3. **(D)** Once this MT switches to the “bright” state, the “dim” lattice progressively switches at the boundary towards the seed, until the MT is entirely “bright”. 40 nM mGFP-EB3. **(E)** Mean intensities of mGFP-EB3 binding to MTs for each mGFP-EB3 concentration, error bars represent standard deviations. The same data are replotted in Fig. 3C. **(F)** Stacked bar chart of MT growth behavior at different mGFP-EB3 concentrations. mGFP-EB3 binding patterns were assessed manually by comparing to the other MTs in the same field of view. “Dim” and “bright” MTs showed relatively uniform binding throughout growth. Some MT segments could not be classified into one of the above categories. *N* indicates the number of MTs considered for analysis from two different experimental replica, (three for 20 nM mGFP-EB3). All experiments were performed with 12.5 μM E254N tubulin. Scale bars 10 μm for length and 5 min for time.

Increasing the EB3 concentration affected the stability of the two conformational lattice states. While an overall average increase of the mGFP-EB3 fluorescence along the E254N MTs was observed as the EB3 concentration was increased from 5 to 40 nM (Fig. 2E), segments with higher and lower EB3 binding density were still present at all concentrations studied. At higher EB3 concentrations, more strongly EB3 binding segments and fewer weakly EB3 binding segments were observed (Fig. 2F). Moreover, with increasing EB3 concentration, slow lattice transformations at the boundary between two segments from weak to high EB3 binding affinity became more frequent. This demonstrates that EB3 promotes the formation of its high affinity binding conformation of E254N MTs.

mGFP-EB3 fluorescence line profiles and intensity histograms demonstrated that the segmented strong and weak mGFP-EB3 binding to E254N MTs was clearly different from the previously described, more uniform high affinity mGFP-EB3 binding to E254A MTs (21) (Fig. 3A, B). Particularly, the intensity histograms show the appearance of a second, high affinity EB3 binding state for E254N MTs with increasing EB3 concentration (Fig. 3B right), explaining why EB3 binding increases more than linearly with increasing EB3 concentration in the tested concentration range, in contrast to the less variable EB3 binding to E254A MTs (Fig. 3C). Plotting the standard deviation of mGFP-EB3 intensity against the mGFP-EB3 intensity also confirms that E254A MTs display a less heterogenous lattice state with respect to EB3 binding than E254N MTs, although some EB3 binding variation is also observable for E254A MTs (Fig. 3D).

**Fig. 3.**
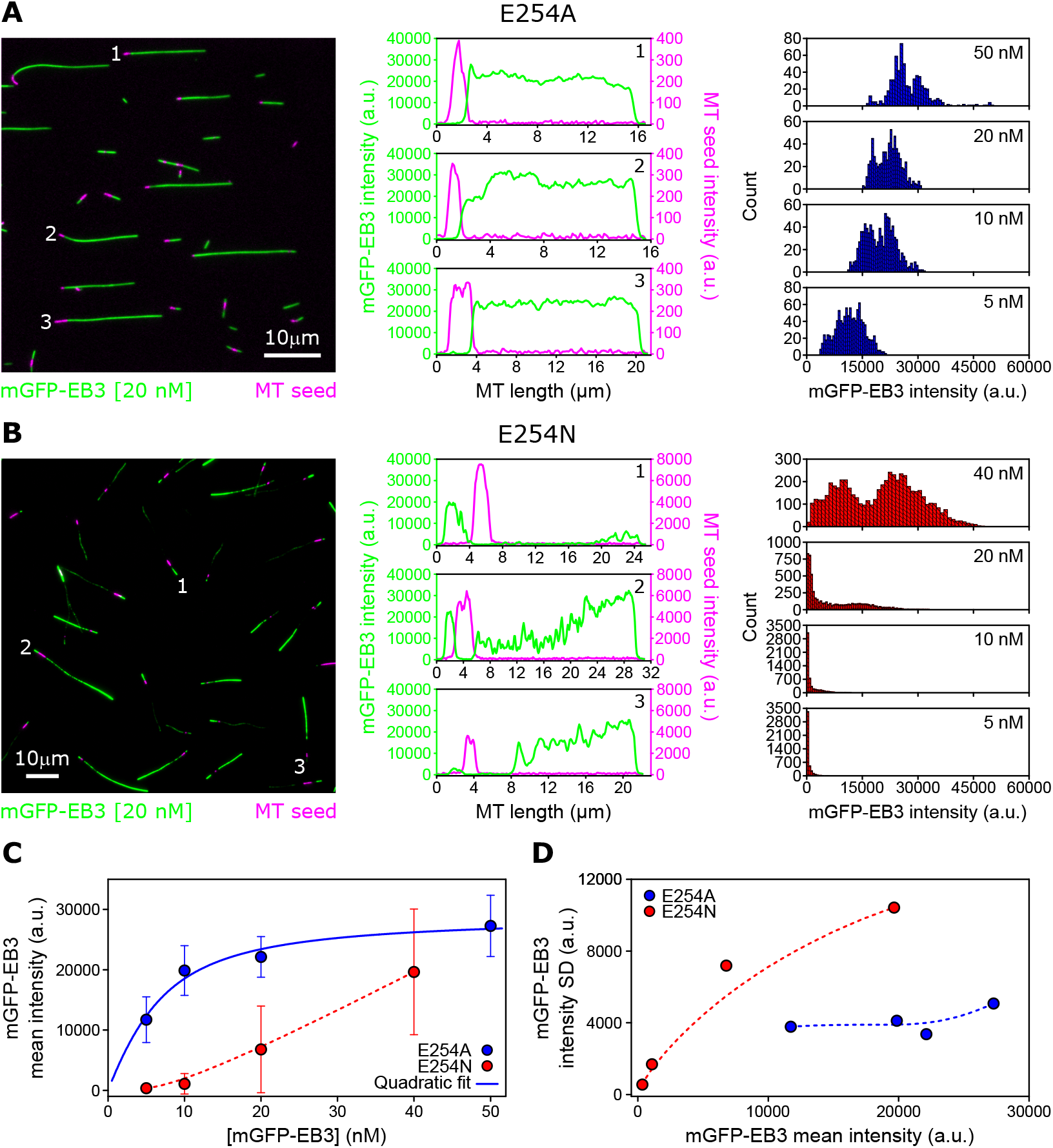
Quantitative comparison of mGFP-EB3 binding to E254A and E254N MTs as observed by TIRF microscopy. **(A-B)** Left: TIRF microscopy images of non-fluorescent E254A MTs **(A)** and E254N MTs **(B)** grown from CF640R-labeled GMPCPP-stabilized “seeds” (magenta) in the presence of 20 nM mGFP-EB3 (green) and GTP. Scale bars represent 10 μm. Middle: line profiles of mGFP-EB3 (green) and MT seed (magenta) intensities along the three MTs indicated in the images on the left. Right: global mGFP-EB3 intensity distribution along several MTs at different mGFP-EB3 concentrations (A-mutant: 5 nM mGFP-EB3, *N* = 18 MTs; 10 nM mGFP-EB3, *N* = 13 MTs; 20 nM mGFP-EB3, *N*= 10 MTs; 50 nM mGFP-EB3, *N* = 11 MTs. N-mutant: 5 nM mGFP-EB3, *N* = 35 MTs; 10 nM mGFP-EB3, *N* = 37 MTs; 20 nM mGFP-EB3, *N* = 43 MTs; 40 nM mGFP-EB3, *N* = 38 MTs). **(C)**Mean intensity of mGFP-EB3 along E254A MTs (blue circles) and E254N MTs (red circles, same data as in Fig. 2E) as a function of mGFP-EB3 concentration. Error bars represent standard deviations. The solid blue line shows a quadratic fit through the E254A MT data, the red dashed line is a Bezier interpolation used as guide-to-the-eye. **(D)** Standard deviation of mGFP-EB3 intensity along the MTs as a function of mGFP-EB3 mean intensity for E254A MTs (blue circles) and E254N MTs (red circles). Dashed lines are Bezier interpolations used as guide-to-the-eye.

These TIRF microscopy experiments show that the two GTP hydrolysis-deficient mutant MTs studied here both can adopt a high affinity binding conformation for EBs. However, E254N MT additionally display a low affinity conformation that can switch into the high affinity conformation during growth, a switch that is promoted by EB binding.

### E254N MTs have an expanded lattice with a positive pf twist in the absence of EB3

Next, we used cryo-EM to determine the lattice parameters of E254N MTs. In contrast to E254A MTs, we did not detect a prominent population of 13 pf 4-start MT species in the E254N MTs. The dominant species of 13pf 3-start E254N MTs showed an expanded dimer rise of 83.45 Å, similar to E254A MTs. However, in contrast to E254A MTs, the pf twist was positive with an angle of +0.13° (Data Table 1). The lattice parameters of E254N MTs are similar to GMPCPP-MTs, where the lattice has a rise of 83.95Å, and a twist of +0.23°. Furthermore, comparison of backbone deviations for the tubulin dimer between the E254N MTs and either GMPCPP-MTs or GDP-MTs clearly shows the E254N and GMPCPP similarity (Fig. S3). This indicates that GTP-MTs in the absence of EBs tend to have an expanded MT lattice, but mutant hydrolysis-deficient GTP-MTs can display different pf twists depending on the particular mutant, revealing an unexpected plasticity of the GTP lattice.

Because the weakly EB3 binding E254N lattice conformation dominated at low EB3 concentrations in our TIRF microscopy experiments, we assign the cryo-EM structure obtained in the absence of EB3 to the weakly EB3 binding lattice conformation observed in TIRF microscopy. This is also in agreement with GMPCPP-MTs, which bind EB3 only very weakly and display similar lattice parameters as E254N MTs in the absence of EB3 (9).

### EB3 compacts both E254N and E254A MTs

Previous cryo-EM work has shown that EB3 has the ability to promote MT compaction and even induce GMPCPP hydrolysis in GMPCPP-MTs (8). Here we observed by TIRF microscopy that EB3 can induce a conformational change in GTP containing E254N MTs revealed by a change in EB binding affinity (Fig. 2, 3). We therefore sought to see what effect EB3 could have at a structural level on the mutant MTs that were locked in a GTP-bound state. Addition of saturating concentrations of the MT binding domain of EB3 to preformed E254N MTs caused the MT dimer rise to decrease from 83.45Å to 81.15Å, indicative of a lattice compaction toward the GDP-like conformation (Data Table 1, Fig. 4A, B). Interestingly, the structure of the EB3-bound catalytically inactive E254N MT clearly shows that the GTP present in the undecorated mutant MTs (Fig. 4C) is still present in the active site, even in the compacted lattice (Fig. 4D). A compacted lattice has also been observed for the γ-phosphate containing GTPγS-EB3 MT lattice (8). The pf twist of - 0.29° agrees well with previous structures of MTs decorated with EB3 that also showed a pf twist of around −0.3° (8, 9). Therefore, we assigned this cryo-EM structure to the strongly EB3 binding lattice conformation of E254N MTs observed by TIRF microscopy.

**Fig. 4.**
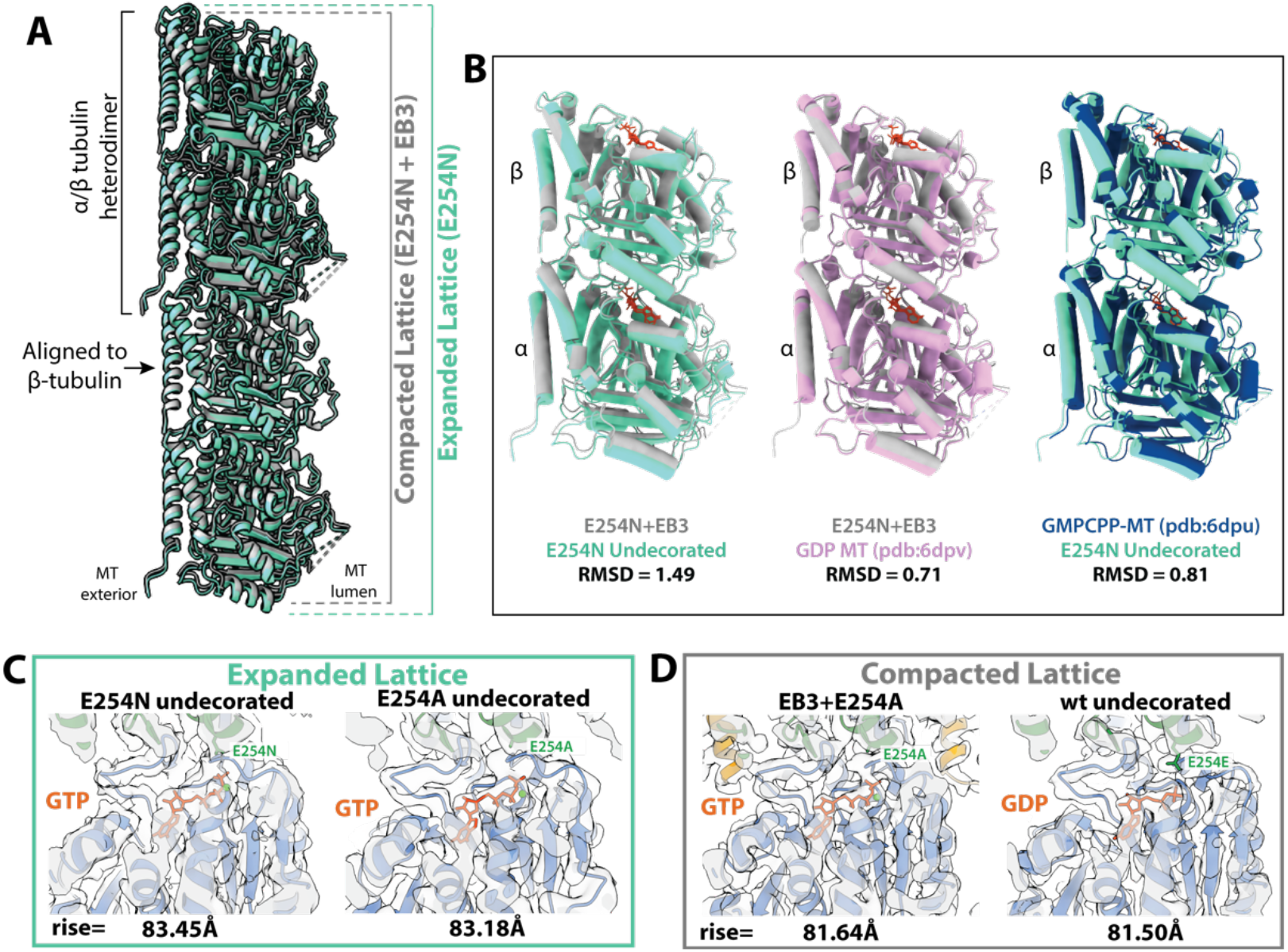
GTP state and compaction for wildtype, E254N, and E254A MTs. **(A)** Visualization of how the MT axial repeat changes, either through hydrolysis of GTP or through EB3 binding to catalytically inactive MTs. Specifically, the panel shows atomic models for the structures determined herein, with the undecorated E254N MT shown in green, and E254N+EB3 MT in grey. **(B)** Comparison of tubulin heterodimers to show the structural changes that occur upon compaction. Dimers are aligned onto β-tubulin, which has been shown to undergo the least amount of conformational changes in previous studies and herein. Root mean square deviation values are reported below each comparison. **(C)** Expanded lattices, with a rise >83Å and with clear GTP density. **(D)** Compacted lattices with a rise <82Å can be formed by EB3 binding (while maintaining GTP), or by hydrolysis into the GDP state.

This same compaction phenomenon was observed for the E254A MTs polymerized in the presence of EB3. The EB3 occupancy was higher in this case, likely due to the higher affinity of EB3 binding to the E254A lattice (Fig. 3C). The dimer twist was found to be −0.19° for the 13pf 3-start E254A MTs grown in the presence of EB3 (Data Table 1). Additionally, GTP was still present in the active site, despite the MTs sampling a compacted state again suggests that lattice compaction can occur without hydrolysis regardless of whether EB3 is added before or after MT formation. Overall, these data continue the trend that EB3 compacts MTs and promotes a more negative MT twist.

### The MT seam opens upon GTP hydrolysis

It is generally accepted that during MT growth individual tubulin dimers experience a certain amount of strain to accommodate the lattice architecture (1). This strain is thought to increase after GTP hydrolysis, thus eventually promoting depolymerization. During MT shortening, lateral contacts are lost first, pfs curl outwards, and eventually longitudinal contacts break down. Due to the heterotypic α/β tubulin interactions formed across the seam, the seam has been considered a potential weak point of the MT lattice where “peeling” of pfs could start (8, 9). By calculating displacement values between a symmetrized and a C1 reconstruction (8, 9), we previously found that in mammalian brain GDP-MTs the pfs at the seam break cylindrical symmetry and appear separated with respect to the rest (we see the same for recombinant, wildtype MTs; Fig. 5A), while this characteristic is either missing or less pronounced in the case of MTs containing non-hydrolyzable GTP analogs (8, 9). The presence of EB3, which binds between pfs, regularizes the seam, irrespective of nucleotide content (Fig. 5B). Here we found that E254N and E254A MTs also do not show any pf opening at the seam, either in the presence (Fig. 5C, D) or absence of EB3 (Fig. 5E). These results add further experimental support for the proposal of lateral tubulin interactions at the seam being stronger before GTP hydrolysis, and the seam possibly becoming the weak lateral interface within the MT lattice after GTP hydrolysis (8, 9).

**Fig. 5.**
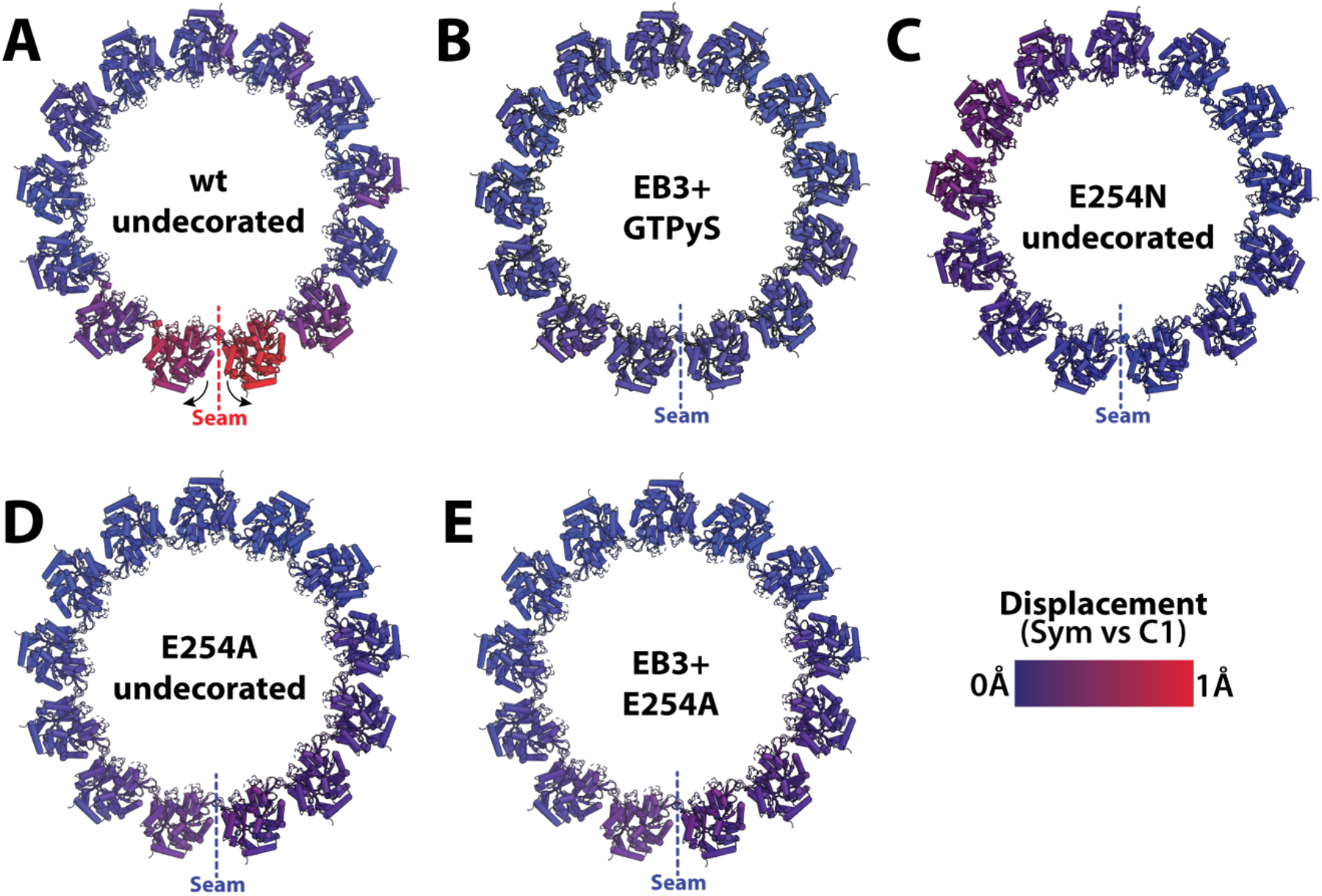
Seam opening correlate with GTP hydrolysis and MT instability. The panels correspond to MT cross-sections for both GTP-like and GDP-MTs showing comparison between the symmetrized and C1 maps to illustrate the brake of symmetry with the opening between pfs at the seam for some of the MTs studied. **(A)** Wildtype MTs in a post-hydrolysis state shows the seam opening phenomenon localized to the pfs at the seam. Movie S1 allows for a dynamic visualization of this motion. (**B**) GTPγS co-polymerized with EB3 originally from data collected for the map deposited as EMD-6347 (8). **(C)** Undecorated E254N-MTs, and **(D)** undecorated E254A MTs. **(E)** The E254A-MT co-polymerized with EB3. No seam opening is seen for B, C, D, or E.

## Discussion

Here we have determined the first high-resolution cryo-EM structures of MTs with GTP bound at the exchangeable nucleotide binding site by visualizing GTP hydrolysis-deficient MTs generated by polymerizing recombinant human tubulin with either a E254A or E254N substitution in α-tubulin. This approach is complementary to previous studies of wildtype MTs grown in the presence of different GTP analogs. Both amino acid substitutions in recombinant tubulin produced MTs lacking catastrophes, as expected for GTP-MTs. Structural analysis revealed that both E254A and E254N MTs display an expanded MT lattice, as was previously described for GMPCPP-MTs (5, 31), but not for GTPγS or BeF_3_^-^ MTs (8, 11). On the other hand, the lattices of the two types of mutant GTP-MTs were not identical, with the major difference concerning pf twist.

When growing as a 13 pf 3-start lattice, which is the lattice that is typically observed in human cells, E254A MTs displayed a negative pf twist, in contrast to the positive twist of 13 pf 3-start E254N MTs. This makes E254N MTs more similar structurally to GMPCPP-MTs. Our observations with mutant, GTP hydrolysis-deficient MTs and previous observations with MTs grown in the presence of slowly hydrolysable GTP analogs demonstrate a remarkable plasticity of the GTP(-like) MT lattice. Furthermore, small changes at the longitudinal tubulin-tubulin interface, either in the nucleotide itself or in the normally catalytic residue, can have a profound impact on MT structure, probably because it is exactly this part of the MT that is designed to be highly sensitive to the small change resulting from GTP hydrolysis. The fact that the tubulin residues critical for hydrolysis also provide interactions at the polymerization interface makes mutated catalytically inactive MTs sensitive to specific changes of this residue.

Regarding the 13 pf 4-start lattice observed for the E254A mutant, different start numbers for MTs have been characterized previously, and although start switching was rare for 13pf MTs, it has been observed (30). Perhaps the conditions used in our cryo-EM studies allowed the E254A MTs to display a variety of lattice structures with respect to pfs and start number—possibly a consequence of high stability of these MTs (21). Of note, the 13pf 4-start phenomenon has also been observed recently in a GDP-BeF_3_^-^ analog (11), but was not analyzed further.

The unexpected lattice plasticity of these MTs may indicate increased flexibility of the GTP-bound state that could play a functional role during tubulin incorporation into the MT and its recognition by cellular factors. However, the 13pf 4-start structure could also be due to the single point mutation introduced in E254A α-tubulin. It is possible that an artificial pf tilting could be caused by the cavity-creating alanine mutation at the inter-dimer interface (Fig. 1A), although, taken together with the observation of this lattice in the GDP-BeF_3_^-^ analog MTs, this structural feature may not be a coincidence.

Another interesting question is what is the lattice structure recognized by EBs? MTs with high EB binding affinity all have a lattice with a negative pf twist, even in the absence of EBs (GTP-E254A, GTPγS-wt, BeF_3_^-^ -wt) (9, 11, 22). MTs to which EBs only bind weakly show either minimal pf twist of ~0.1Å (GDP-wt) or they show a positive twist (GMPCPP-wt, GTP-E254N) (8, 9). Moreover, when EB3 is forced to bind to lattices with a positive pf twist (GMPCPP-wt, GTP-E254N) under saturating conditions, the pf twist becomes negative, suggesting that the high affinity conformation of the EB binding site at the corners of 4 tubulins requires a negative pf twist (8). Additionally, EB3 binding at saturating conditions compacts normally extended MT lattices (GMPCPP-wt, GTP-E254N, GTP-E254A), irrespective of whether the nucleotide can be hydrolyzed (GMPCPP-wt) or not (GTP-E254N, GTP-E254A) (9). The compaction of the GTP lattice may be somewhat inhibited in E254N MTs, explaining their segmented appearance, while it occurs more easily in E254A MTs, explaining the higher affinity of EB binding to E254A MTs. Given that EBs have the ability to compact GTP-bound MTs and to accelerate a conformational transition towards the GDP lattice (32), it is likely that compaction occurs before hydrolysis in wildtype MTs growing in the presence of GTP, a model recently proposed based on studies of multiple GTP analogs (11).

Taken all this information together, we favor a model for the structure of the GTP cap in wildtype MTs in which the GTP lattice transitions through several conformations before forming a mature GDP lattice (Fig. 6). Initially, freshly formed pfs would be in an expanded state, possibly sampling conformations corresponding to what it is observed in GTP-E254A, GTP-E254N, GMPCPP-MTs. Lattices with a positive twist show weak EB binding affinity and may contribute to the very short zone at the very tip of growing MT ends showing only weak EB binding (21). However, such weakly EB binding lattices are likely to be only short-lived; otherwise, there would be a longer EB-undecorated part before the “EB comet” starts at the growing end of wildtype MTs. The high affinity EB binding region likely has a negative twist, possibly transitioning from an expanded GTP-E254A-like lattice to a compacted GTPγS-wt-like lattice, in agreement with recent evidence suggesting that the GTPγS lattice also corresponds to a GTP state (11). How fast this transition takes place, remains unknown, because both lattices show high EB binding affinity. Hydrolysis followed by phosphate release would then cause the pf twist to become ~0° in the GDP lattice to which EBs only bind weakly. EBs may already bind weakly to the part of the MT that still contains the phosphate after hydrolysis and before release. A recent cryo-EM structure with trapped GDP and phosphate in the presence of the MT-stabilizing protein doublecortin displayed a compacted lattice with only little twist to which EB3 is not expected to bind well (10).

**Fig. 6:**
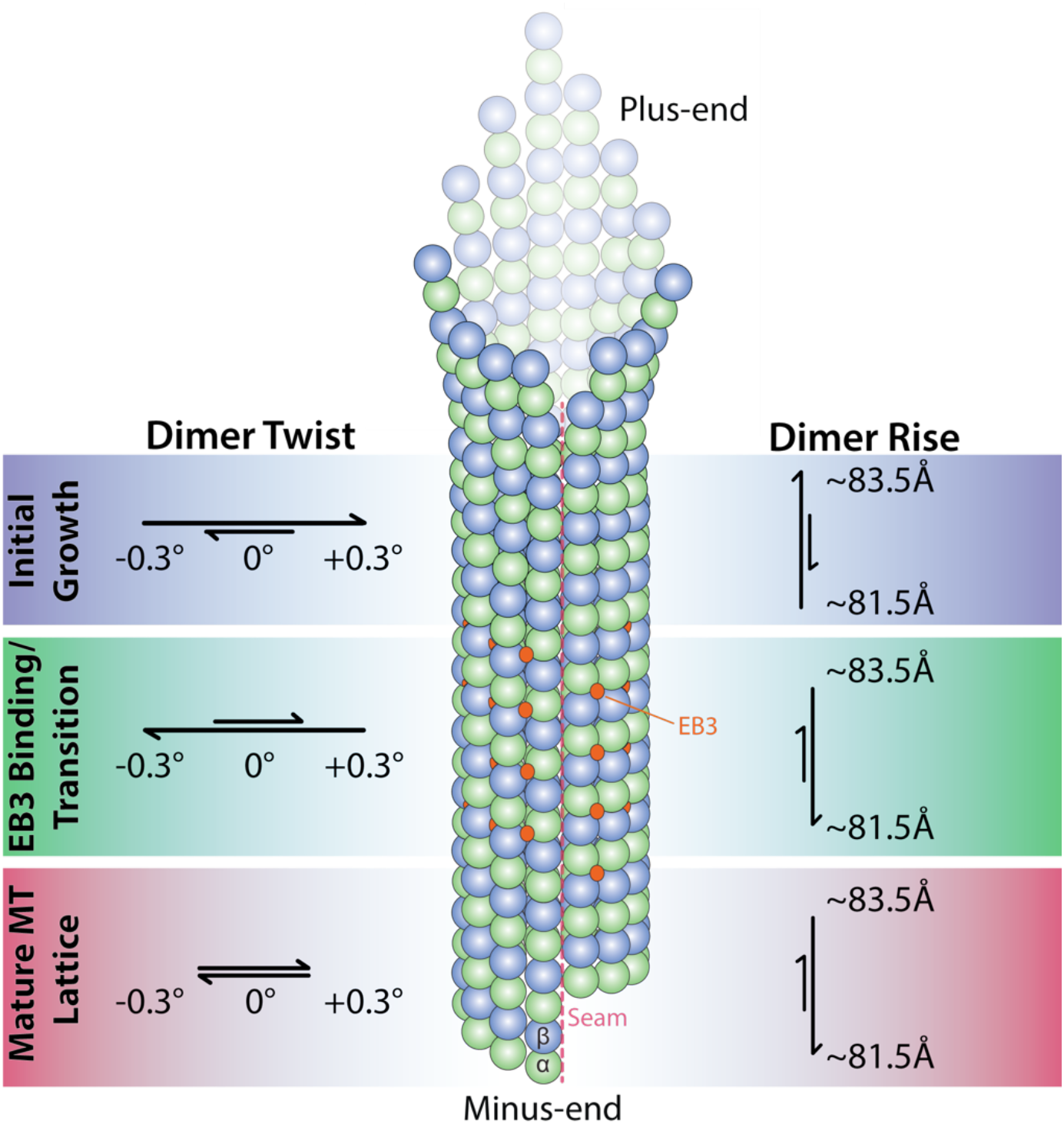
Model of MT growth informed by cryo-EM and TIRF microscopy observations

The exact interactions within the MT lattice that confer greater stability in the GTP lattice relative to the GDP lattice remain unclear, at least in part because no high-resolution structure of a true GTP-wildtype lattice is available. We have previously proposed that the tension present in the GDP-lattice can be directly visualized as a seam-opening phenomenon, which could have the greatest effect at the GTP-to-GDP transition zone. This model was put forward considering that GMPCPP gives rise to a GTP-like MT lattice. Our present study, using hydrolysis mutants instead of GTP analogs, further supports such a model, as the stable hydrolysis-inactive MTs all exhibit a “closed” seam structure, as do GMPCPP and GTPγS MTs, as well as MTs stabilized by drugs (8, 9, 33).

Thus, MT seam stability may serve as a proxy for overall MT stability. The more symmetric pf arrangement around the MT axis observed for the E254N and E254A mutant (Fig. 5 C, D), and for GMPCPP-MTs (8, 9), is associated with the more stable GTP region of MTs. On the other hand, the symmetry-breaking, outward rotation of the seam-proximal pfs in GDP-MTs both for the wildtype MTs described here (Fig. 5 E), and in the previously described porcine MTs (9), reflects the tension at the seam upon hydrolysis that likely precedes pf peeling and depolymerization when exposed at a MT end.

## Conclusions

Recent advances in the production of recombinant tubulin have opened up the door to new structure-function studies of MTs. In this work, we utilized recombinant human tubulin to better understand the effect of GTP hydrolysis on MT structure and stability, and thus the mechanistic basis of the property of dynamic instability essential for MT cellular function. We show that hydrolysis-impaired E254A and E254N mutant MTs display an expanded lattice conformation and a seam lacking the structural sign of strain seen in GDP-MTs that have undergone nucleotide hydrolysis (8, 9). Our mutant studies provide additional insights into the conformational dynamics that accompany the GTP-to-GDP transition in MTs, suggesting a series of transitions from an expanded to a compacted lattice with accompanying changes also in pf twist. Future studies visualizing the ends of dynamic MTs with high resolution will ultimately allow us to observe the true GTP cap and test the model for its structure proposed here.

## Materials and Methods

### Purification of recombinant human tubulin

Human tubulin was purified recombinantly as described previously (21). Briefly, cell pellets from 2 liters of High Five insect cells expressing human TUBB3-TEV-Strep and TUBA1B-His^internal^ (wildtype, E254A, or E254D) were resuspended 1:1 (v/v) in cold lysis buffer and lysed by douncing 60 times. Lysate was diluted 4-fold in dilution buffer, and clarified by ultracentrifugation (158,420× g, 1 hr, 4°C). The supernatant was passed through a HisTrap HP column (GE Healthcare), and the elution was immediately diluted 6× in Strep buffer, and passed through a 1mL HiPrep column followed by a 5mL StrepTag HP column. Tubulin was eluted off the column, diluted 2-fold in Strep elution buffer and incubated on ice for 2 hours with TEV protease to remove the Strep-TagII from TUBB3. The eluate was then clarified by ultracentrifugation (204,428× g, 10min, 4°C). The supernatant was further purified through a 1mL HiPrep SP FF column, desalted into storage buffer, concentrated to 3.5mg/mL, ultracentrifuged (278,088× g, 10min, 4°C), and flash frozen in 10uL aliquots with liquid nitrogen.

|  |  |
| --- | --- |
| Lysis | 80mM PIPES, 1mM EGTA, 6mM MgCl<sub>2</sub>, 50mM imidazole, 100mM KCl, 2mM GTP, 1mM BME, pH 7.2 + protease inhibitors and DNase |
| Dilution | 80mM PIPES, 1mM EGTA, 6mM MgCl<sub>2</sub>, 50mM imidazole, 2mM GTP, 1mM BME, pH 7.2 |
| His Elution | 80mM PIPES, 1mM EGTA, 5mM MgCl<sub>2</sub>, 500mM imidazole, 2mM GTP, 1mM BME, pH 7.2 |
| Strep Binding | 80mM PIPES, 1mM EGTA, 5mM MgCl<sub>2</sub>, 2mM GTP, mM BME, pH 7.2 |
| Strep Elution | 80mM PIPES, 1mM EGTA, 4mM MgCl<sub>2</sub>, 2.5mM D-desthiobiotin, 50mM imidazole, 2mM GTP, 1mM BME, pH 7.2 |
| Storage | 80mM PIPES, 1mM EGTA, 1mM MgCl<sub>2</sub>, 0.2mM GTP, pH 6.8 |

### Purification of EB3

Monomeric EB3 used for cryo-EM was purified as previously described (8, 32). Human EB3_1-200_ was inserted into a 2BT vector with a C-terminal HisTag (Macrolab, UC-Berkeley), and expressed in BL21(DE3)-RIL *E. coli*. Cell pellets from a 2L culture were resuspended in 1:2 (v/v) Lysis Buffer, and lysed by sonication. Cell debris was pelleted by centrifugation (18,000× g, 45min, 4°C), and supernatant was loaded on a 5mL HisTrap column (GE Healthcare), and eluted with a 0-100% gradient of Lysis buffer to elution buffer. The elution was incubated with TEV protease overnight at 4°C to remove the His-tag, followed by a subtractive nickel purification to remove TEV, uncleaved EB3, and the His-tag. The flow-through was concentrated to 500uL and loaded on a Superdex 200 10/300 GL pre-equilibrated in SEC buffer for size-exclusion chromatography. Peak fractions were pooled, concentrated to 20μM, and flash frozen in SEC buffer until needed.

mGFP-EB3 used for TIRF microscopy was purified as described (21).

|  |  |
| --- | --- |
| Lysis | 50mM Tris pH 8.0, 150mM NaCl, 1mM EDTA, 1mM DTT + protease inhibitors, DNase, RNase, and Lysozyme |
| His Elution | 50mM Tris pH 8.0, 150mM NaCl, 1mM EDTA, 1mM DTT, 300mM imidazole |
| SEC Buffer | 50mM Tris pH 8.0, 150mM NaCl, 1mM EDTA, 1mM DTT, 10% glycerol |

### Purification of Kinesin

The kinesin plasmid was generously supplied by the Vale lab (34), and purified as described previously (5). The plasmid encoding His6-tagged, monomeric, catalytically inactive Kif5b (1-350aa, E236A) was transformed into BL21(DE3) cells for expression. Upon reaching OD_600_=0.5, the 1L expression culture was brought to 22°C, and induced with 0.2mM IPTG for 16 hrs. The cells were harvested with by centrifugation at 4000g for 20min at 4°C. Cell pellets were resuspended in Lysis Buffer and incubated at room temperature for 30min followed by sonication (3×45sec, at power level 7). The lysate was clarified by centrifugation at 30,000g for 60min, and the supernatant was then incubated with Nickel-NTA beads (GE Healthcare). The protein was washed with 6 column volumes of wash buffer, and eluted with 2 column volumes of elution buffer. The elution was pooled and loaded onto a size exclusion column equilibrated with SEC buffer. Fractions were analyzed by SDS-PAGE for kinesin, and peak fractions were pooled and concentrated to 20μM before flash freezing in liquid nitrogen until needed.

|  |  |
| --- | --- |
| Lysis | 50mM NaPO<sub>4</sub> pH 8.0, 300mM KCl, 2mM MgCl<sub>2</sub>, 10% glycerol, 10mM Imidazole, 1mM DTT |
| Wash | 50mM NaPO<sub>4</sub> pH 8.0, 150mM KCl, 2mM MgCl<sub>2</sub>, 10% glycerol, 30mM Imidazole, 1mM DTT, 200uM ATP |
| Elution | 50mM NaPO<sub>4</sub> pH 8.0, 150mM KCl, 2mM MgCl<sub>2</sub>, 10% glycerol, 250mM Imidazole, 1mM DTT |
| SEC buffer | 25mM Tris pH 7.5, 150mM KCl, 10% glycerol, 2mM MgCl<sub>2</sub>, 2mM DTT |

### Cryo-EM sample preparation

CryoEM specimens were prepared on CFlat 1.2/1.3-T open hole grids (Protochips) that were plasma cleaned for 30 seconds with a Tergeo plasma cleaner (Pie Scientific). A 10μL aliquot of tubulin was thawed on ice for 10min and supplemented with 0.05% NP-40. The tubulin was incubated at 37°C for 30-40min to form MTs. While polymerizing, the Vitrobot Mark IV was equilibrated to the following conditions: 37°C, 100% humidity, 15 blot force, and a 4 second blot with 1 second drain time. 2μL of MT solution was adsorbed onto the grid for 30 seconds, followed by plunge freezing into a eutectic solution of liquid ethane/propane (70:30). For conditions where Kinesin or EB3 were used to decorate the MT, two 4μL aliquots of 20μM kinesin/EB3 were added to the MT grid with a 30 second wait time in between additions to allow binding. Blotting conditions were identical for all mutants and decorating MT associated proteins to maintain consistency.

### Cryo-EM Data Collection

All grids were clipped and loaded into a Gatan Autoloader for imaging at the BACEM Titan Krios microscope at UC-Berkeley. The data was collected with a GIF energy filter and Gatan K2 or K3 camera (depending on the dataset) operating in super-resolution mode. Data collection was controlled by SerialEM (35). Each micrograph had a total electron exposure of 40e^-^ collected over 40 frames and was collected within a defocus range of 0.8-2.0μm. Collection parameters for each dataset are reported in Data Table S1. The E254A structure was obtained over 2 different data collection sessions because a more thorough classification was necessary. It should be noted that the same grid was used for both sessions, and there was no bias in the number of the 13pf 4-start particles for either of the data collections.

### Cryo-EM Analysis and Model Building

Data processing was done mostly within the RELION framework (36). Final processing steps that are specific to MTs were performed in FREALIGN which was necessary for identifying the correct seam location for undecorated MTs (37). Briefly, MotionCorr 2.1 5×5 patch-based alignment was performed on each micrograph (38). CTFFind4 was used to estimate the defocus of each micrograph (39). MTs were manually picked within RELION, and extracted with a repeat length of 82Å. Initial pf classification was performed with RELION’s Class3D function on bin4 data. Once the helical parameters were classified, particles within the same class (primarily 13 pf) were unbinned, re-centered, and re-extracted. Unbinned particles were refined within RELION, and then converted into FREALIGN format for pseudo-helical Fourier symmetrization to improve resolution, as well as SeamSearch protocols (29) to correctly identify the seam location. Data processing procedures are summarized in Fig. S2.

Atomic models were built using a previous porcine MT structure as a template (pdb: 6dpu), changing the necessary residues to account for the differences between porcine and human tubulins as well as the active site mutations. Each tubulin subunit was rigid-body docked with phenix, and refined using the real space refinement program within PHENIX (40, 41). All refinements were treated with the same number of iterations in order to minimize variations in the processing procedure.

For the seam analysis (Fig. 5), atomic coordinates were rigid-body docked into both the C1 and pseudo-helical symmetry refined maps. The displacement was calculated using the *colorbyrmsd.py* script within PyMol (PyMOL Molecular Graphics System, Schrödinger LLC) and then normalized so that all samples were on the same scale from 0 to 1 Å. *colorbyrmsd.py* was also used to calculate displacements between various MT states in Fig. S3.

### Determination of the MT Nucleotide Content

The nucleotide content of E254N MTs (Fig. S1B) was determined by HPLC as described (21).

### TIRF Microscopy Assay with Recombinant Human Tubulin Mutants

Dynamic MT assays with E254N tubulin were performed using a TIRF microscope (Cairn Research, Faversham, UK), as described previously (21, 42). In brief, flow chambers were prepared from poly-(L-lysine)-polyethylene glycol (PEG) passivated microscopy slides and biotin-PEG-functionalized coverslips (43). Chambers were further passivated for 5 minutes at room temperature with 5% Pluronic F-127 (P2443, Sigma-Aldrich) and extensively washed with assay buffer (80 mM PIPES, 1 mM EGTA, 1 mM MgCl_2_, 30 mM KCl, 1 mM GTP, 5 mM 2-ME, 0.15% (w/vol) methylcellulose (4000 cP, Sigma-Aldrich), 1% (w/vol) glucose, pH 6.8), followed by 2 washes with κ-casein (50 μg/ml in assay buffer, C0406, Sigma-Aldrich) in assay buffer. Neutravidin diluted in the κ-casein solution (50 μg/ml, A2666, Thermofisher) was flowed in, incubated for 3 minutes at room temperature, washed out with assay buffer, followed by biotinylated GMPCPP porcine brain MT seeds labelled with CF640R (12% labelling ratio) diluted in assay buffer. After a final wash with assay buffer, the final reaction mix was flowed into the flow chamber. This mix included 98% unlabeled recombinant human E254N tubulin diluted in assay buffer containing oxygen scavengers (180 mg/ml catalase (C40, Merck) and 752 mg/ml glucose oxidase (22778.01, Serva)) and 2% mGFP-EB3 diluted in its storage buffer (50 mM Na-phosphate, 400 mM KCl, 5 mM MgCl_2_, 0.5 mM 2-ME, pH 7.2) to final concentrations of 12.5 mM E254N tubulin and 5-40 nM mGFP-EB3. Directly after sealing, the flow chamber was placed inside the microscope incubator at 30°C and imaged after 2-3 minutes with a 100X oil-immersion objective (Nikon CFI SR Apo, NA = 1.49) for 20 minutes at 1 frame/5 s with 300 ms exposure times for both MT seeds (using 640 nm laser excitation) and mGFP-EB3 (using 480 nm laser excitation). 25 minutes afterwards the initial temperature shift, several distinct fields of view were acquired for the quantification of mGFP-EB3 intensity profiles along the MTs.

Data with E254A MTs were recorded previously as described (21). In brief, E254A MTs were polymerized from (non-immobilized) fluorescent, biotinylated GMPCPP-stabilized porcine brain MT seeds (0.5 μM polymerized tubulin) at 1 μM E254A tubulin for 1 hr at 37°C in BRB80 (80 mM PIPES, 1 mM EGTA, 1 mM MgCl_2_, pH 6.8) with 1 mM GTP. These stable MTs were stored at room temperature until use. Flow chambers were prepared as above, except that κ-casein was added to the chamber placed on a metal block on ice. Prepolymerized, stable MTs diluted in BRB80 were attached to the neutravidin surface at room temperature. The chambers were then washed and imaged using 98% assay buffer with oxygen scavengers as above and supplemented with 2% mGFP-EB3 diluted in its storage buffer, yielding a final mGFP-EB3 concentration between 5 and 50 nM.

### TIRF Microscopy Imaging

Dynamic TIRF microscopy assays with E254N tubulin and mGFP-EB3 were performed using a total internal reflection fluorescence (TIRF) microscope (Cairn Research, Faversham, UK) (42). Directly after sealing the flow chamber, it was placed inside the incubator at 30°C and after 2-3 minutes imaged with a 100x oil-immersion objective (Nikon CFI SR Apo, NA = 1.49) for 20 minutes at 1 frame/5 s with 300 ms exposure times for both MT seeds (using 640 nm laser excitation) and mGFP-EB3 (using 480 nm laser excitation). Several images in different fields of view were acquired 25 minutes after the initial temperature shift for the measurements of mGFP intensity profiles along the MTs. The data for stable E254A MTs with mGFP-EB3 are from a previous study (21) and were re-analyzed in detail here.

### TIRF Microscopy Data Analysis

All images were processed using FIJI (v1.53c, RRID:SCR_002285) (44). Images were first corrected for uneven illumination using the FIJI rolling ball background subtraction algorithm with a radius of 50 pixels. Then, segmented lines (3 pixel wide) were drawn along the longer MT segment elongating from the seed, excluding overlapping MTs. Line intensity profiles were averaged over their width. For each intensity profile, the local background was obtained by shifting the segmented line to the closest region without MTs and subtracted pixel-by-pixel. Background-subtracted intensity profiles were pooled together to generate mGFP-EB3 global intensity distributions and calculate their moments using OriginPro2021 (OriginLab, RRID:SCR_014212).

Kymographs from individual E245E MTs were obtained using custom FIJI macros to trace maximum-intensity projections. The different mGFP-EB3 binding patterns to E245N MTs were categorized and quantified manually.

## Author Contributions

BJL designed experiments, collected, processed, and interpreted data for the cryo-EM portion of the project, and wrote the initial draft of the manuscript. JR designed and produced the recombinant tubulin mutants and performed HPLC and TIRF microscopy experiments. GH performed TIRF microscopy experiments, and analyzed TIRF microscopy data. BJG aided in analysis and interpretation of cryo-EM data. RZ and CM performed initial cryo-EM experiments. DN analyzed TIRF microscopy data. TS and EN guided research and edited the manuscript. All authors edited and contributed to the final version of the paper.

## Acknowledgments

We thank Claire Thomas for help with expressing and purifying recombinant tubulin in insect cells, Abhiram Chintangal and Paul Tobias for support with computation, Dan Toso, Jonathan Remis, and Patricia Grob for support with electron microscopy, Simone Kunzelmann and Iain Taylor who helped with the determination of nucleotide content of E254N MTs by HPLC and Juan Estevez Gallego for his critical reading of the manuscript. BJL was supported by an NSF-GRFP grant 1106400. JR, GH and TS were supported by the Francis Crick Institute, which receives its core funding from Cancer Research UK (FC001163), the UK Medical Research Council (FC001163), and the Wellcome Trust (FC001163). JR was supported by a Sir Henry Wellcome Postdoctoral Fellowship (100145/Z/12/Z). EN acknowledges support from the National Institutes of Health (R35GM127018). TS acknowledges support from the European Research Council (Advanced Grant, project 323042). GH, DN and TS acknowledge the support of the Spanish Ministry of Economy, Industry and Competitiveness to the CRG-EMBL partnership, the Centro de Excelencia Severo Ochoa and the CERCA Programme of the Generalitat de Catalunya. TS also acknowledges support from the Miller Institute for Basic Research in Science at UC Berkeley. EN is a Howard Hughes Medical Institute Investigator.

---

**Data Table 1.** Lattice parameters for select 13pf MT structures.

| Sample | Lattice Spacing/<br>Dimer Rise (Å) | Skew/<br>Twist (°) | Intradimer<br>Rise (Å) | Interdimer<br>Rise (Å) | Reference |
| --- | --- | --- | --- | --- | --- |
| GMPCPP-TPX2 | 84.33 | +0.40 | 41.67 | 42.61 | Zhang et al, 2018 |
| GMPCPP | 83.95 | +0.23 | 41.53 | 42.43 | Zhang et al, 2018 |
| GMPCPP-DCX | 83.74 | +0.16 | NA | NA | Manka et al, 2018b |
| GMPCPP-Kinesin | 83.16 | +0.19 | 41.45 | 42.00 | Zhang et al, 2018 |
| **E254N** | **83.45** | **+0.13** | **41.53** | **41.93** | **This Study** |
| **E254A 3-start** | **83.18** | **-0.22** | **41.15** | **42.36** | **This Study** |
| **E254A 4-start** | **83.03** | **+2.40** | **NA** | **NA** | **This Study** |
| GDP-P<sub>i</sub>-DCX | 81.74 | 0.00 | NA | NA | Manka et al, 2018b |
| GTPγS-Kinesin-EB3 | 81.96 | -0.28 | 41.31 | 40.51 | Zhang et al, 2018 |
| GTPγS | 81.93 | -0.12 | 41.35 | 40.45 | Zhang et al, 2018 |
| GTPγS-EB3 | 81.52 | -0.25 | 41.30 | 40.30 | Zhang et al, 2015 |
| GTPγS-DCX | 82.10 | -0.02 | NA | NA | Manka et al, 2018b |
| GDP-DCX | 81.90 | +0.08 | NA | NA | Manka et al, 2018b |
| GDP | 81.76 | +0.08 | 41.44 | 40.30 | Zhang et al, 2018 |
| **wt E254-GDP-Kinesin** | **81.69** | **-0.09** | **41.38** | **40.37** | **This Study** |
| **EB3+E254A** | **81.64** | **-0.19** | **41.25** | **40.44** | **This Study** |
| GDP-Kinesin | 81.64 | +0.01 | 41.38 | 40.36 | Zhang et al, 2018 |
| GDP-Kinesin-EB3 | 81.59 | -0.11 | 41.35 | 40.32 | Zhang et al, 2018 |
| **wt E254-GDP** | **81.50** | **+0.05** | **41.25** | **40.30** | **This Study** |
| GDP-EB3 | 81.44 | -0.12 | 41.40 | 40.30 | Zhang et al, 2015 |
| **E254N+EB3** | **81.15** | **-0.29** | **41.35** | **40.45** | **This Study** |

MT lattice parameters obtained in this study are reported alongside previously published values for reference. Expanded lattices are in blue (GMPCPP and undecorated catalytically inactive mutants); presumed transition state MTs (GDP+P<sub>i</sub> or GTPγS) are colored in green, and compacted lattices are shaded in red (GDP state, or catalytically inactive mutants + EB3). NA signifies cases where intra-dimer or inter-dimer values were not reported due to mixing of $\alpha/\beta$ -tubulin subunits, or where resolution was too low to accurately determine the values.

