## Supplemental Figures for "Structural transitions in the GTP cap visualized by cryo-EM of catalytically inactive microtubules"

(A)

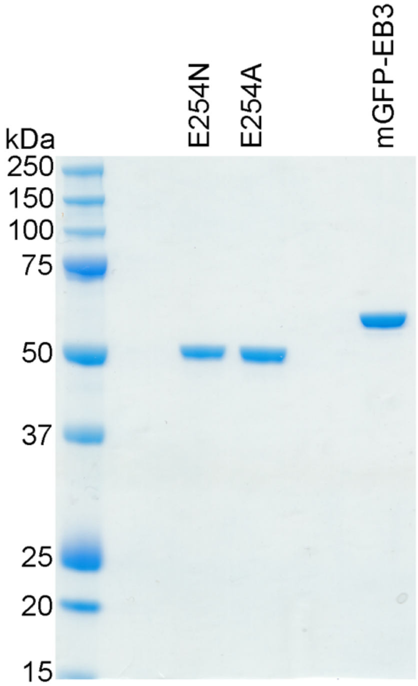


(B)

**E254N nucleotide content**

| | % GDP | % GTP |
|---|---|---|
| **exp. 1** | 12.1 | 87.9 |
| **exp. 2** | 16.6 | 83.4 |
| **exp. 3** | 14.9 | 85.1 |
| **Average** | **14.5** | **85.5** |
| **STDEV** | **2.3** | **2.3** |

**Figure S1. Biochemical characterization of mutant tubulin.**
**(A)** Coomassie-stained SDS gel showing E254N tubulin, E254A tubulin, and mGFP-EB3 used for TIRF microscopy. **(B)** Table showing the detected GDP and GTP content of E254N MTs as obtained in three independent experiments by HPLC. The detected GTP content is similar to that of E254A MTs (1).

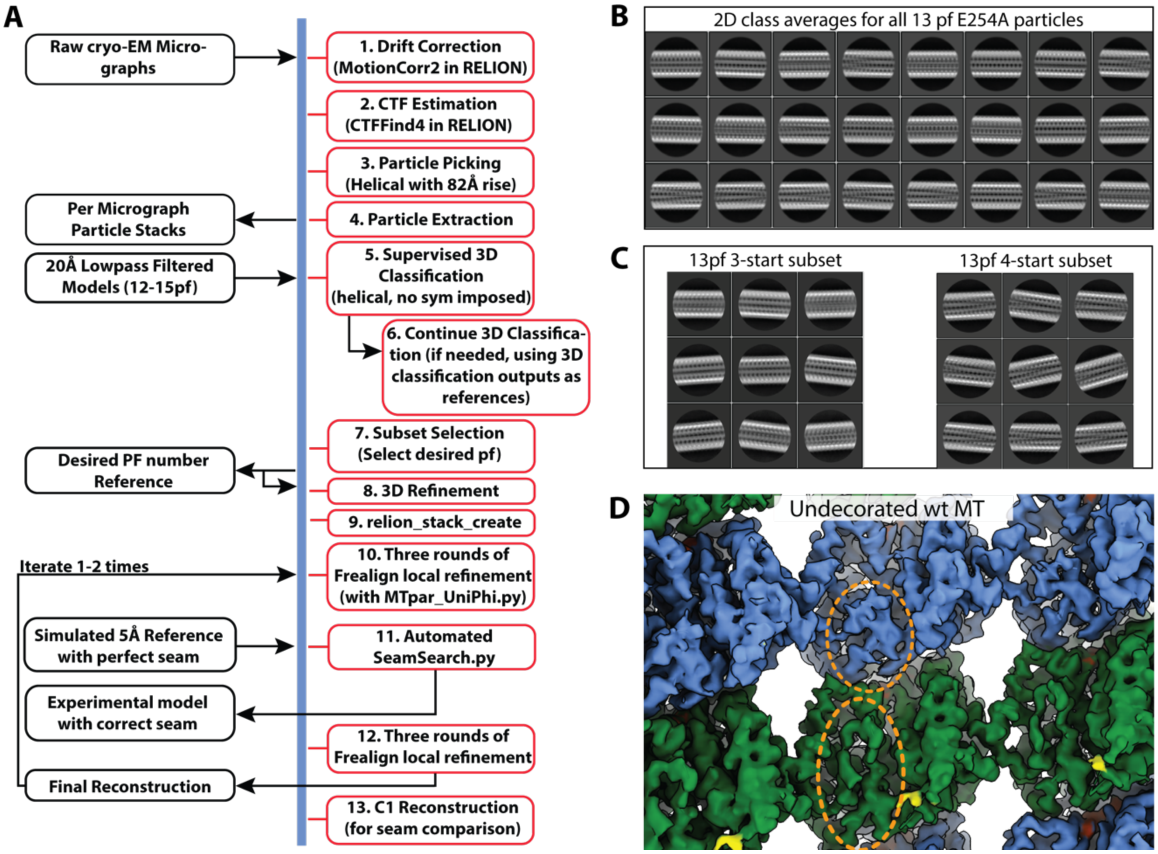


**Figure S2. Cryo-EM data processing pipeline with example classes and structure.**
**(A)** Schematic of the data processing pipeline used to reconstruct MT structures based on a hybrid approach between the recently described MIRP protocol (2) and the SeamSearch technique (3). Steps 1-9 are performed within the RELION framework. In order to resolve the 3-start and 4-start structures for E254A, the output from the first round of supervised 3D classification, along with the original references, were used to subclassify 13pf models. This was done 3 times, until no residual classification was observed. Furthermore, because the dataset corresponding to undecorated MTs (without an associated protein such as EB3 to serve as fiducial for the tubulin dimer), SeamSearch was necessary to separate α- and β-tubulin, as outlined in steps 10-12. **(B)** Initial 2D classification results obtained from the E254A dataset. **(C)** After classification based on protofilament type, an additional sub-classification for 13pf MTs to separate dimer twist revealed both “straight” classes (pfs running parallel to the MT axis in the 3-start lattice, left) and “super twisted” classes (corresponding to the 4-start lattice, right). **(D)** Structure of wildtype recombinant MTs and separation of α- and β-tubulin for undecorated MTs. The small region highlighted in yellow corresponds to additional density in the recombinant MTs that can be assigned to the internal $His_6$-tag present in β-tubulin. The distinction of α- and β-tubulin subunits (green and blue, respectively) obtained with our image analysis scheme in the absence of decoration with other factors is highlighted by the dashed orange ovals marking a distinctly longer loop in α-tubulin facing the MT lumen.

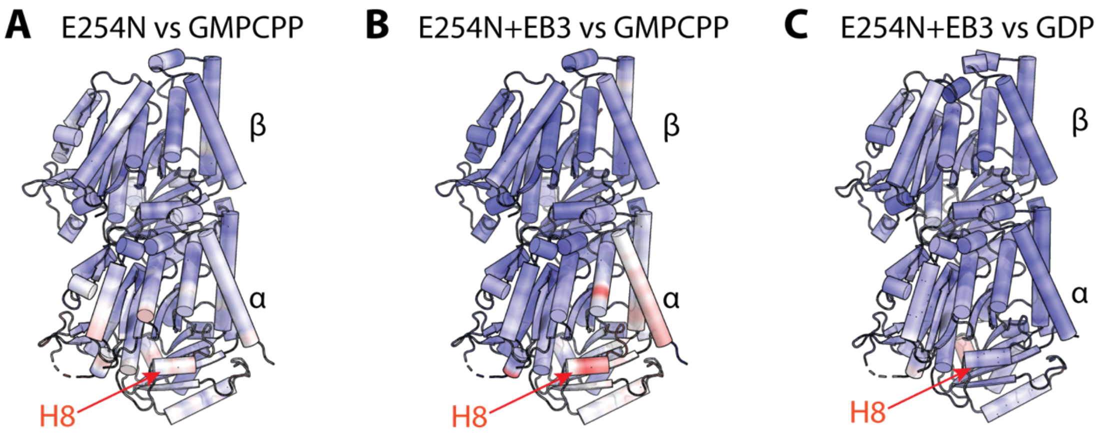


**Figure S3. Mutant dimer structure versus the GMPCPP and GDP states within the MT.** Displacement values for the tubulin dimer from various MT structures after alignment to β-tubulin and displayed after normalization to the same scale for all models (see Materials and Methods, blue-to-red coloring indicates 0Å-1Å displacement, respectively). The location of helix 8 (H8) which has been previously shown to have the greatest displacement up compaction (4) is indicated by the red arrow in each panel. This figure shows that in addition to the lattice parameters reported in Data Table 1, E254N resembles the GMPCPP state at the tubulin dimer structural level (A). Once bound by EB3, and accompanying lattice compaction, the EB3+E245N tubulin structure deviates from the GMPCPP MT structure (B) and very closely resembles that seen in the GDP MT (C).

**Table S1. Data collection, 3D reconstruction, and refinement statistics.**

| **Dataset** | **wt** | **wt + kinesin** | **E254A (3/4)** | **EB3+E254A** | **E254N** | **E254N+EB3** |
|---|---|---|---|---|---|---|
| Microscope | Titan Krios | Titan Krios | Titan Krios | Titan Krios | Titan Krios | Titan Krios |
| Stage type | Autoloader | Autoloader | Autoloader | Autoloader | Autoloader | Autoloader |
| Voltage (kV) | 300 | 300 | 300 | 300 | 300 | 300 |
| Detector | Gatan K2 | Gatan K2 | Gatan K3 | Gatan K3 | Gatan K3 | Gatan K3 |
| Data Collection Software | SerialEM | SerialEM | SerialEM | SerialEM | SerialEM | SerialEM |
| Acquisition mode | Super-res | Super-res | Super-res | Super-res | Super-res | Super-res |
| Physical pixel size (Å) | 0.575 | 0.460 | 0.595 | 0.460 | 0.575 | 0.575 |
| Defocus range (μm) | 0.5-2.5 | 0.7-2.4 | 0.6-2.5 | 0.7-2.5 | 1.0-2.7 | 1.0-2.7 |
| Electron exposure ($e^{-}/A^{2}$) | 40 | 40 | 40 | 40 | 40 | 40 |
| **Reconstruction** | **EMD-XXXX** | **EMD-XXXX** | **EMD-XXXX** | **EMD-XXXX** | **EMD-XXXX** | **EMD-XXXX** |
| Session | 18Dec07b | 18Dec07c | 19Jun03 | 20Sep10 | 20Feb03 | 20Feb04 |
| Software | RELION 3.1 & Frealign | RELION 3.1 & Frealign | RELION 3.1 & Frealign | RELION 3.1 & Frealign | RELION 3.1 & Frealign | RELION 3.1 & Frealign |
| Particles picked | 33,575 | 39,703 | 165,039 | 77,608 | 77,703 | 6,465 |
| Particles final (13pf) | 19,365 | 23,264 | 3-start: 68000<br>4-start: 26022 | 56,705 | 13,706 | 3,825 |
| Extraction box size (pixels) | $512^3$ | $512^3$ | $512^3$ | $512^3$ | $512^3$ | $256^3$ |
| Final pixel size (Å) | 0.92 | 0.92 | 1.19 | 0.92 | 1.15 | 2.30 |
| Map resolution (Sym; Å) | 3.8 | 3.6 | 3-start: 3.4<br>4-start: 3.7 | 3.5 | 3.8 | 5.0 |
| Map sharpening B-factor ($Å^2$) | -88 | -92 | 3-start: -38<br>4-start: -50 | -72 | -36 | -100 |
| **Coordinate refinement** | | | | | | |
| Software | PHENIX | PHENIX | -- | PHENIX | PHENIX | -- |
| Refinement algorithm | REAL SPACE | REAL SPACE | -- | REAL SPACE | REAL SPACE | -- |
| Resolution cutoff (Å) | 3.8 | 3.8 | -- | 3.7 | 3.8 | -- |
| $FSC_{model-vs-map}$=0.5 (Å) | 3.9 | 4.0 | -- | 3.6 | 4.2 | -- |
| **Model** | **PDB-XXXX** | **PDB-XXXX** | **--** | **PDB-XXXX** | **PDB-XXXX** | **--** |
| Number of residues | 864 | 864 | -- | 995 | 861 | -- |
| B-factor overall | 115 | 92 | -- | 144 | 120 | -- |
| R.m.s. deviations | | | | | | |
| Bond lengths (Å) | 0.006 | 0.005 | -- | 0.003 | 0.004 | -- |
| Bond angles (°) | 0.606 | 0.563 | -- | 0.632 | 0.545 | -- |
| **Validation** | | | | | | |
| Molprobity clashscore | 12.57 | 11.81 | -- | 14.11 | 13.43 | -- |
| Rotamer outliers (%) | 5.4 | 5.9 | -- | 1.0 | 5.8 | -- |
| $C_\beta$ deviations (%) | 0.0 | 0.0 | -- | 0.0 | 0.0 | -- |
| Ramachandran plot | | | | | | |
| Favored (%) | 95.2 | 95.7 | -- | 96.6 | 95.0 | -- |
| Allowed (%) | 4.8 | 4.3 | -- | 3.4 | 5.0 | -- |
| Outliers (%) | 0.0 | 0.0 | -- | 0.0 | 0.0 | -- |

This table notes the microscope collection parameters, as well as the map and model values used for all the final reconstructions and atomic models.

**Movie S1 (separate file).** Morph between E254N MTs and wildtype MTs showing the rotation of pfs proximal to the seam.
